# Mechanistic characterization of a *Drosophila* model of paraneoplastic nephrotic syndrome

**DOI:** 10.1101/2023.04.23.538006

**Authors:** Jun Xu, Ying Liu, Weihang Chen, Joshua Shing Shun Li, Aram Comjean, Yanhui Hu, Norbert Perrimon

## Abstract

Paraneoplastic syndromes occur in cancer patients and originate from dysfunction of organs at a distance from the tumor or its metastasis. A wide range of organs can be affected in paraneoplastic syndromes; however, the pathological mechanisms by which tumors influence host organs are poorly understood. Recent studies in the fly uncovered that tumor secreted factors target host organs, leading to pathological effects. In this study, using a *Drosophila* gut tumor model, we characterized a mechanism of tumor-induced kidney dysfunction. Specifically, we found that Pvf1, a PDGF/VEGF signaling ligand, secreted by gut tumors activates the PvR/JNK/Jra signaling pathway in the principal cells of the kidney, leading to mis-expression of renal genes and paraneoplastic renal syndrome-like phenotypes. Our study describes a novel mechanism by which gut tumors perturb the function of the kidney, which might be of clinical relevance for the treatment of paraneoplastic syndromes.

## Introduction

Paraneoplastic syndromes are a group of disorders associated with tumors that are prevalent in patients with lymphatic, lung, ovarian or breast cancers^1, 2^. These syndromes are not attributable to tumor invasion or compression but are due to the response of body systems and organs to tumors and/or the factors that they secrete^3, 4^. As such, characterization of tumor-host interactions might help reveal mechanisms underlying paraneoplastic syndromes. However, due to the complexity of tumor-host interactions in patients and in mammalian cancer models, studies in alternative and simpler models of the mechanisms underlying paraneoplastic syndrome pathogenesis are needed.

A wide range of organs can be affected by cancer, and dysfunction in different body systems leads to distinct paraneoplastic syndromes^3^. For instance, dysfunctions in the nervous system caused remotely by tumors, termed paraneoplastic neurologic disorders, lead to a diverse group of symptoms such as encephalitis, optic neuropathy, and cerebellar degeneration^5^. In other cases, polymyositis and hypertrophic osteoarthropathy have been reported as paraneoplastic syndromes caused by influence of lung cancer on muscle and bone, respectively^6, 7^. Defects in the kidney have also been observed in patients with cancer. However, in these instances, it is not clear whether the renal dysfunctions are caused by treatment of the cancer or whether tumor-secreted factors such as hormones and cytokines are the primary cause^8–10^.

*Drosophila* has emerged as a simple and powerful model to identity pathogenic tumor-host interactions^11, 12^. The fly has organs that are functionally equivalent to human organs, including digestive tract, liver, adipose tissue, and kidney^13^, and a number of models have been established to study organ-specific disorders triggered by tumors^11, 12^. For instance, elevated JAK-STAT signaling in a fly tumor model disrupts the blood-brain barrier leading to death^14^. In addition, two tumor-secreted factors, Unpaired 3 (Upd3) and PDGF- and VEGF-related factor 1 (Pvf1), target host muscle and fat body to stimulate body wasting, which is reminiscent of cachexia^15, 16^. Here, leveraging a *Drosophila* gut tumor model, we demonstrate that tumor-secreted PDGF- and VEGF-related factor 1 (Pvf1) stimulates PDGF/VEGF signaling in the fly malpighian tubules (MTs), a renal system functionally equivalent to the vertebrate kidney^17, 18^, leading to formation of kidney stones, ascites/bloating, and uric acid accumulation. Importantly, these renal phenotypes contribute to increased mortality of flies with gut tumors. Altogether, our data suggests that tumors remotely induce renal dysfunction through hijacking of normal signaling pathways, which represents an underappreciated effect of paraneoplastic renal syndromes. Our findings that a tumor can induce renal disorders, independent of cancer treatment, might open new therapeutic avenues to treat kidney-related symptoms in patients with cancer.

## Results

### Kidney dysfunction contributes to bloating and reduces lifespan of Yki flies

Flies with gut tumors induced by activated *yorkie* (*esg> yki^act^*; referred to hereafter as Yki flies) exhibit a ‘bloating phenotype’ characterized by an enlarged, fluid-filled abdomen^19^. This suggests the possibility of a defect in the fly renal system, the Malpighian tubules (MTs), which maintain fluid and electrolyte balance in the fly hemolymph^20^. Interestingly, the MTs of Yki flies contain crystals (kidney stones) in their lower segments, which increase in size upon tumor progression (Fig. 1A). The accumulation of kidney stones can be driven by many factors, including elevated levels of uric acid^21, 22^. Indeed, uric acid levels are increased in Yki flies as compared to controls (Fig. 1B). To assess the role of uric acid levels in the formation of kidney stones, we fed Yki flies with allopurinol, which can repress uric acid levels by inhibiting purine degradation^21^. Although wild type flies are tolerant, this drug led to severe lethality of Yki flies. As an alternative strategy, we increased uric acid levels by feeding Yki flies with high purine food, which induced kidney stones in the main segment of the MTs (Fig. 1CD), indicating that uric acid levels positively correlate with kidney stone formation. To determine if formation of kidney stones impacts renal function, we fed Yki flies with 3% *Garcinia cambogia* extract that is known to dissolve kidney stones in flies^23^, and observed a reduction in average kidney stone sizes from 18.12um to 4.67um without affecting the gut tumors (Fig. 1E). Interestingly, *Garcinia cambogia* feeding significantly decreased the prevalence of bloating among tumor flies from 86.4% to 35.5% (Fig. 1FG). Consistent with this, the average wet body weight of tumor flies was reduced from 2.52 mg to 1.54 mg, while the dry mass remained unchanged (Fig. 1H). Yki flies fed with *Garcinia cambogia* did not show a reduction in uric acid levels (Fig. S1A), likely because hydroxycitric acid dissolves kidney stones by inhibiting calcium oxalate monohydrate nucleation^24^. Next, to determine whether kidney stones lead to reduced lifespan, we fed flies with 0.5% calcium oxalate (NaOx), which induces stone formation in the lumen of MTs^25^. NaOx-fed flies showed an increase in kidney stone size and reduced lifespan (50% mortality was shifted from 18 days to 5 days) (Fig. 1IJ). Altogether, these data suggest that kidney disfunction contributes to the imbalance of body fluids and mortality of Yki flies.

**Figure 1.**
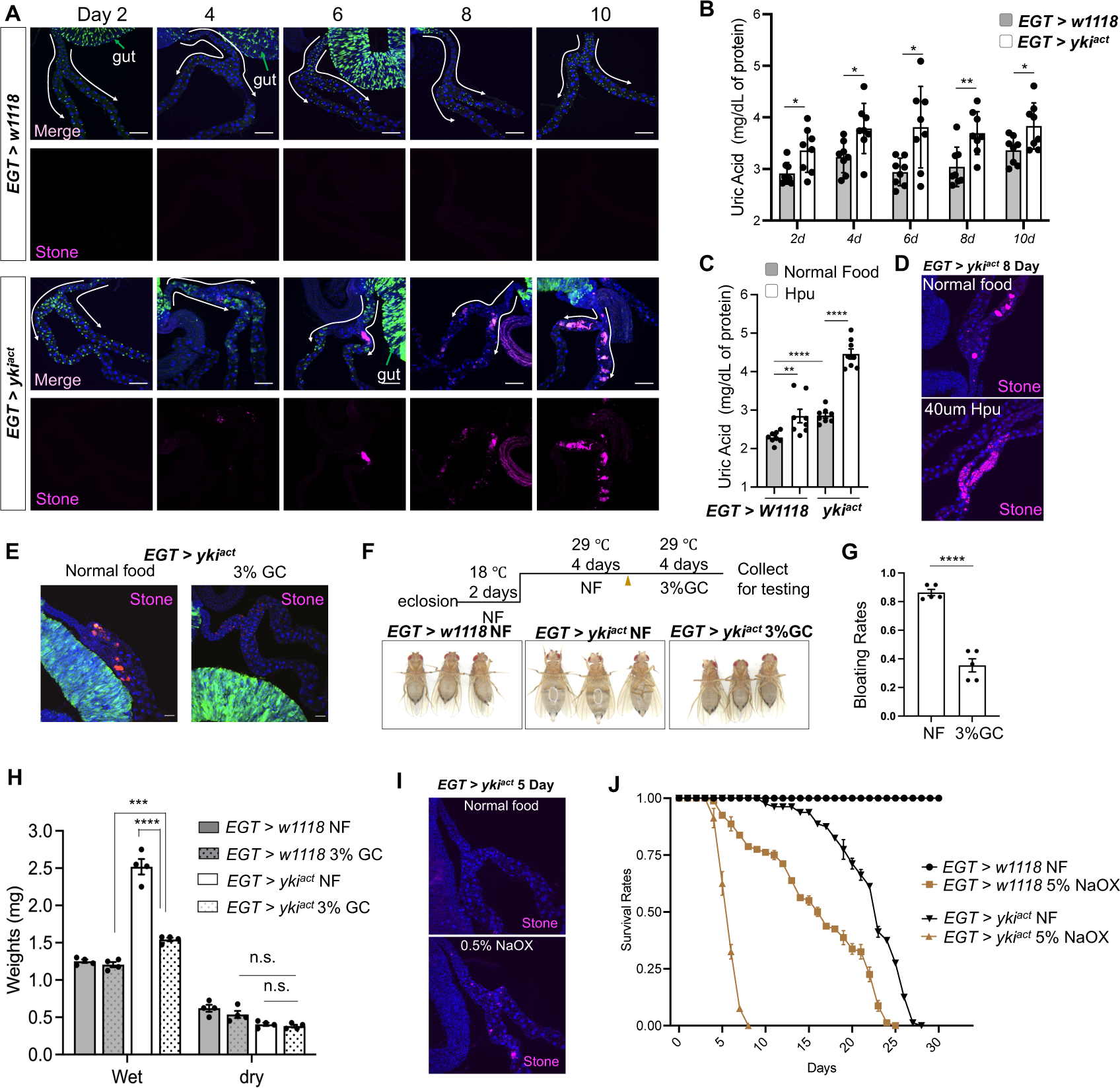
*EGT > yki^act^*gut tumor triggers kidney stone formation and uric acid (UA) accumulation. (A) Changes in kidney stone appearance (purple) in the MT of *EGT > yki^act^*flies (*esg-GAL4, UAS-GFP, tub-GAL80^TS^ /+; UAS-yki^3SA^/+)* and *EGT > w1118* control (*esg-GAL4, UAS-GFP, tub-GAL80^TS^ /+*) 2 days to 10 days after tumor induction. Green shows *esg* positive cells, blue is DAPI staining for nuclei. White arrows indicate the renal stem cell zone of MT, green arrows show the gut. (B) Whole-body UA level in *EGT > w1118* and *EGT > yki^act^* (N=8) flies at different time points. (C-D) High purine food (Hpu) feeding increases the amount of (C) whole body UA levels and (D) kidney stones. (E) Dissolution of kidney stones in *EGT > yki^act^*flies upon feeding *Garcinia cambogia* (GC) and (F-G) inhibition of the bloating phenotype. (H) Wet/dry weights showing that GC feeding affects the water volume in the body. Feeding with sodium oxalate (NaOx) (I) increased kidney stones and (J) decreased lifespan of flies. Data are presented as means ± SEM. *p < 0.05, **p < 0.01, ***p < 0.001, ****p < 0.0001. n.s. means no significant with student t-test.

### Single-cell survey reveals abnormal cell composition of Malpighian tubules with Yki gut tumors

To gain a comprehensive understanding of renal dysfunction in Yki flies, we performed a single nuclei RNA-sequencing (snRNA-seq) analysis of MTs from control and Yki flies. We analyzed MTs at day 8 upon tumor induction when the bloating phenotype is apparent^15^. In total, 1706 and 1849 cells from control and Yki flies, respectively, were recovered from snRNA-seq. Cell clusters were annotated according to our previous report of marker genes in MTs (Fig. 2A)^26^. Nine cell clusters were identified, including initial and transitional principal cells (PCs), main segment PCs, lower tubule PCs, upper ureter PCs, lower ureter PCs, lower segment PCs, renal stem cells (RSCs), and stellate cells (SCs) (Fig. 2A). An additional cluster was identified in the Yki sample, which we labelled as ‘main segment PCs 2’ as they express *List* (Fig. 2A), a previously identified main segment PC marker gene^26^. To assess changes in MTs in response to Yki gut tumors, we analyzed the respective number of cells within each cluster. Yki fly MTs displayed a decrease in main segment PCs and an increase in upper ureter PCs and lower segment PCs (Fig. 2B), suggesting abnormalities in these renal segments.

**Figure 2.**
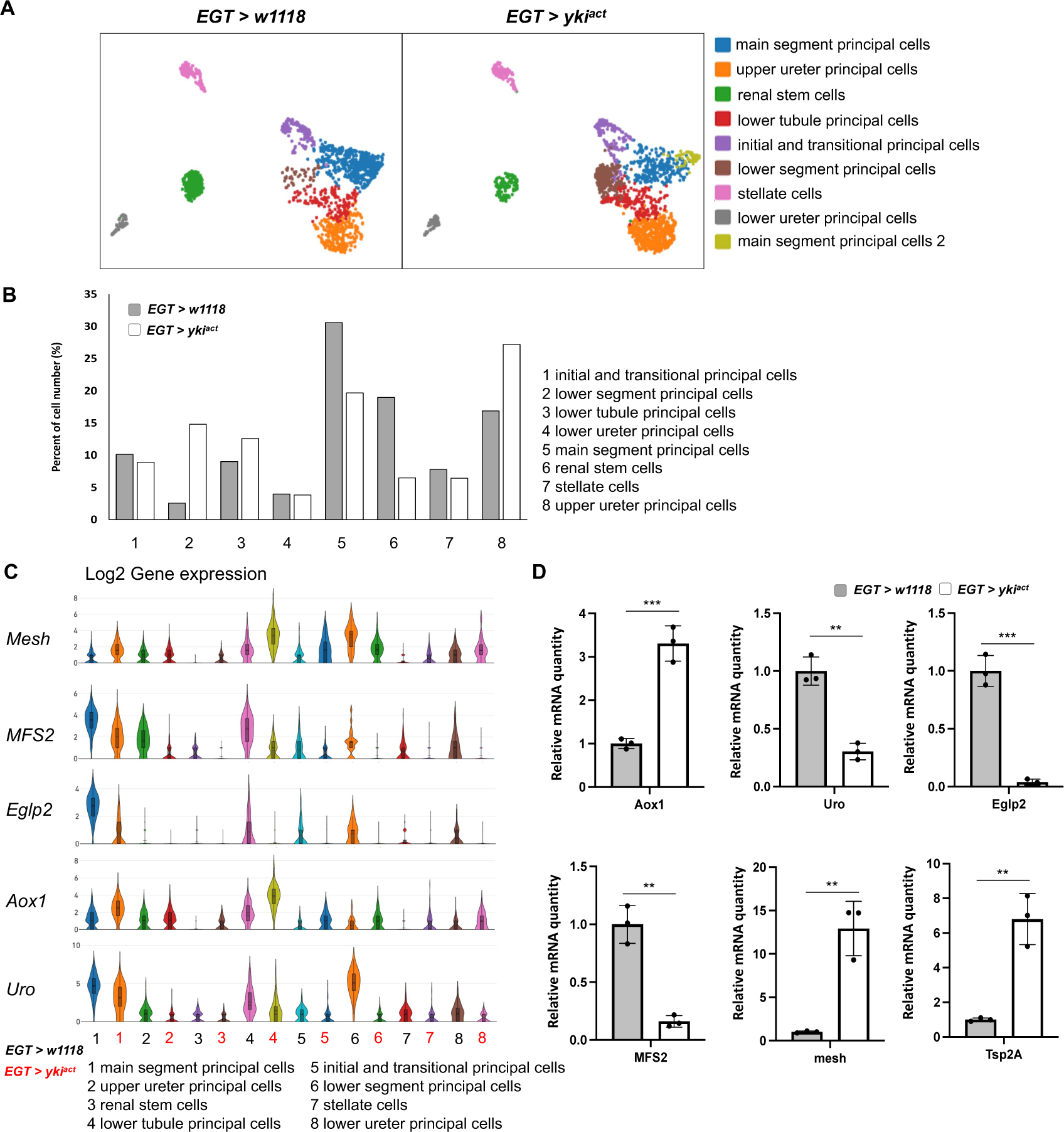
Gene expression in MT of *EGT > yki^act^*flies. (A) A UMAP indicating the MT cell types of *EGT > w1118* and *EGT > yki^act^* flies. Cell clusters corresponding to principal cells (initial and transitional principal cells, main segment principal cells, lower tubule principal cells, upper ureter principal cells, lower ureter principal cells, and lower segment principal cells), renal stem cells and stellate cells. (B) Percentage of cells in each cluster in *EGT > w1118* and *EGT > yki^act^* flies. (C) Violin plots of the expression change in the different cell clusters of genes involved in various renal functions in *EGT > w1118* and *EGT > yki^act^* flies. (D) qPCR analysis indicating the changes in gene expression levels in the MT of *EGT > w1118* and *EGT > yki^act^* flies. Transgene expression was induced for 8 days. Data are presented as means ± SEM. *p < 0.05, **p < 0.01, ***p < 0.001 with student t-test.

Next, we analyzed the expression levels of genes important for kidney function, which include genes involved in cell junction, kidney stone formation, cation transport, diuretic, aquaporin, and uric acid metabolism (Fig. 2CD, Fig. S2A-I). Consistent with the elevated uric acid levels in Yki flies, the expression of two uric acid pathway genes, *Uro* and *CG30016*, which promote uric acid excretion^21, 26^, was significantly down-regulated. Conversely, we observed an increase in expression in Yki flies of *Aldehyde oxidase 1* (*AOX1*) (Fig. 2CD, Fig. S2G, I), which catalyzes xanthine into uric acid^27^. In addition, kidney stone disease-related genes such as *CG31674, Vacuolar H+-ATPase 55kD subunit* (*Vha55*), and *Major Facilitator Superfamily Transporter 2* (*MFS2*) were all down-regulated in Yki flies (Fig. 2CD, Fig. S2C, I). Previous studies have shown that depletion of *Vha55* or *MFS2* in PCs can induce kidney stone formation^23, 28^. As such, the reduced levels of these genes likely contribute to kidney stone formation in Yki flies. Further, to investigate the impaired water balance in Yki flies, we assessed the expression of genes involved in cation transport, diuresis, and water transport. Cation transport genes, including *MFS2* and *Prestin,* were down-regulated (Fig. 2CD, Fig. S2C, I). In addition, the *Secretory chloride channel* (*SecCl*) (2.44, p = 0.0054) was up-regulated, suggesting an impaired response to diuretic hormones (Fig. S2D, I)^29^. Moreover, genes encoding Aquaporins, *Entomoglyceroporin 2* (*Eglp2*) (0.039, p = 0.00025), *Eglp4* (0.19, p = 0.00018), and *Prip* (0.24, p = 0.00097), were all down-regulated (Fig. 2CD, Fig. S2F, I). Changes in expression of these genes presumably reduce the ability of MTs to transport water, leading to excessive buildup of body fluids.

Also, genes involved in cell junctions, including *Innexin 7* (*Inx7*) (1.95, p = 0.025), *Inx2* (1.74, p = 0.029), *Tetraspanin 2A* (*Tsp2A*) (6.80, p = 0.0024), *mesh* (12.93, p = 0.0027), *Snakeskin* (*Ssk*) (3.65, p = 0.00033) and *discs large 1* (*dlg1*) (1.66, p = 0.034) were up-regulated in Yki flies (Fig. 2CD, Fig. S2B, I), suggesting a dysregulation of renal structure. Altogether, these results indicate that key renal genes are mis-expressed in Yki flies, leading to impairment in renal functions (Fig. S2H).

### Elevated PDGF/VEGF signaling in principal cells causes kidney dysfunction

Yki tumors secrete Pvf1 and Upd3, which activate the Pvr/MEK and JAK/STAT signaling pathways, respectively, and ImpL2, which represses insulin/insulin-like signaling (IIS), in peripheral tissues^15, 16, 19^. Thus, we tested whether any of these pathways were involved in the renal dysfunction observed in Yki flies. Since genes involved in kidney stone formation and uric acid metabolism are highly enriched in PCs (Fig. S2C and G), we first searched for Gal4 drivers that would allow us to manipulate the activity of these pathways in a PC-specific manner. The two widely used MT drivers, *C42-Gal4* and *Uro-Gal4*, are either not PC-specific or are weakly expressed. Thus, we analyzed genes preferentially expressed in PCs for candidate PC-specific Gal4 drivers. Among them, *CG31272* showed strong and specific expression in all PCs, which was confirmed with a Gal4 splicing trap (CG31282[MI05026-TG4.1]) (Fig. S3A-D). Combining this Gal4 driver with tub-Gal80ts (hereafter referred to as PC^ts^), we tested activation of PDGF/VEGF and JAK/STAT signaling pathway, and inhibition of IIS in PCs of wildtype flies, mimicking the conditions in Yki flies. Only PDGF/VEGF activation (*Pvr^act^*) in PCs increased whole body uric acid levels (Fig. 3A) and resulted in bloating, increased ratio of water/dry mass, and reduced lifespan, phenocopying what happens in Yki flies (Fig. 3B-D). Importantly, kidney stones were observed when *Pvr^act^*was expressed in MTs (Fig. 3E). In addition, elevated uric acid levels were not observed when PDGF/VEGF was activated in SCs (Fig. S3E), suggesting that SCs are not involved in renal dysfunction in Yki flies. Altogether, these observations indicate that activating the PDGF/VEGF pathway specifically in PCs is sufficient to cause renal dysfunction and phenocopy defects observed in Yki flies.

**Figure 3.**
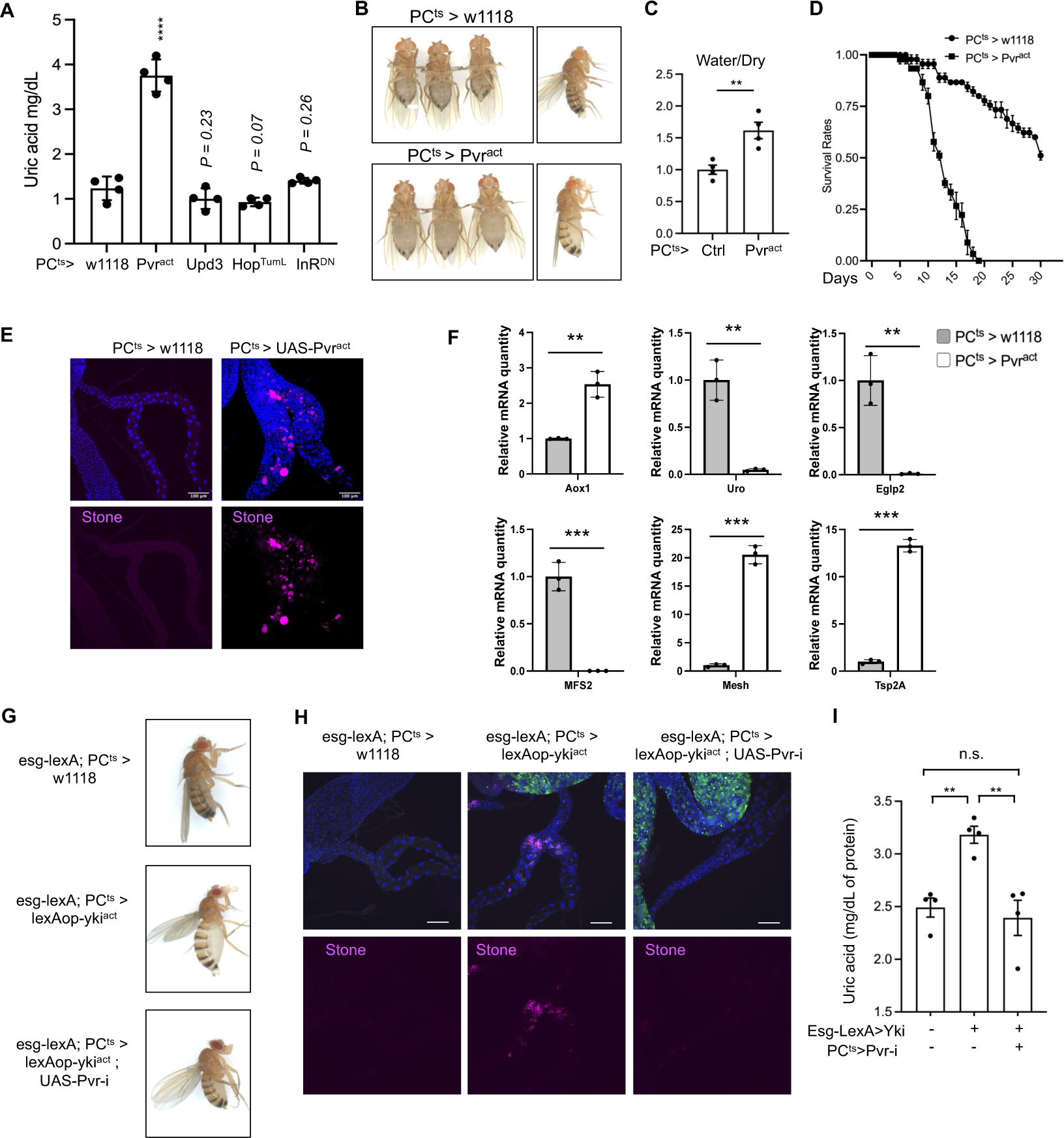
Activation of PDGF/VEGF signaling in principal cells leads to MT dysfunction. (A) Whole-body UA level in *PC^ts^ > Pvr^act^, Upd3*, *Hop^TumL^*, and *InR^DN^* flies (n=4) and *PC^ts^ > w1118* as control (*PC^ts^* corresponds to *tub-GAL80^ts^, CG31272-GAL4*). (B) Bloating phenotype associated with expression of *Pvr^act^* driven by PC^ts^ in principal cells (C) Ratio of fly water/dry mass. (D) Lifespan of *PC^ts^ > Pvr^act^* and *PC^ts^ > w1118* control flies. N=45. (E) Kidney stone images: blue is for DAPI staining to detect nuclei and purple show the kidney stones. (F) qPCR results showing changes in gene expression levels in the MT tubule of *PC^ts^ > Pvr^act^* and *PC^ts^ > w1118* flies. (G-I) Rescue of the bloating, kidney stone and uric acid phenotypes following knockdown of *Pvr* in principal cells in the *EGT > yki^act^* flies. Data are presented as means ± SEM. *p < 0.05, **p < 0.01, ***p < 0.001, ****p < 0.0001. n.s. means no significant with student t-test.

Next, we examined the expression of the genes found to be mis-regulated in Yki flies, in the MT samples with PC-specific activation of PDGF/VEGF signaling (Fig. 3F, Fig.S3F). Changes in expression of these genes were similar to those observed in Yki flies (Fig. 2D and 3F, Fig.S3F). For instance, cell junction genes, including *Inx7* (3.62, p = 0.0006), *Inx2* (3.46, p = 0.0003), *Tsp2A* (13.29, p = 0.0000068), *mesh* (20.55, p = 0.000031), *Ssk* (5.07, p = 0.00032) and *dlg1* (2.66, p = 0.013) were up-regulated (Fig. 3F, Fig. S3E). Further, genes associated with kidney stones, including *CG31674* (0.068, p = 0.00013), *Vha55* (0.25, p = 0.012), *MFS2* (0.0021, p = 0.00033), and *Prestin* (0.36, p = 0.011), were down-regulated, and the Aquaporin genes *Eglp2* (0.011, p = 0.0029) and *Eglp4* (0.0035, p = 0.00054) were also down-regulated (Fig. 3F, Fig. S3E). Uric acid production-related genes *Uro* (0.048, p = 0.00035) and *CG30016* (0.0073, p = 0.000023) were significantly down-regulated, while *AOX1* (2.54, p = 0.0019) was up-regulated (Fig. 3F, Fig. S3E). To further confirm that PVR signaling in MTs of Yki flies is required for the activation of these genes, we used the LexA-LexAop system to induce *yki^act^* tumor in the gut and the Gal4/UAS system to knockdown *Pvr* in PCs. Inhibition of PDGF/VEGF signaling following expression of *Pvr-i* in PCs rescued the bloating, renal stone, and uric acid phenotypes observed in tumor-bearing flies (Fig. 3G-I). Altogether, these results indicate that elevated Pvf1/PDGF/VEGF signaling in tumor flies is responsible for renal dysfunction.

### PDGF/VEGF signaling activates the JNK pathway in principal cells

Pvr signaling is known to activate the Ras/Raf/MAP kinase (ERK) and JNK pathways^30^. This led us to test the roles of the ERK and JNK pathways in the MTs of Yki flies. The JNK pathway regulates the two downstream transcription factors (TFs), *kayak* (*kay*) and *Jun-related antigen* (*Jra*), both at the transcriptional and phosphorylation levels^31–34^. Interestingly, expression of both *kay* and *Jra* was upregulated in Yki flies (Fig. 4AB and E), whereas no change in expression was observed for *anterior open* (*aop),* a TF downstream of ERK signaling (Fig. 4C). Supporting a role for JNK signaling in the MTs of Yki flies, the JNK cascade reporter gene *puckered* (*puc*) was up-regulated in the main segment PCs (Fig. 4D), an observation we confirmed by qRT-PCR (Fig. 4E). In addition, expression of *kay*, *Jra*, and *puc* was also up-regulated in MTs following PC-specific activation of the PDGF/VEGF pathway (Fig. 4F). These results indicate increased JNK pathway activity in MTs of both *Pvr^act^* and Yki flies. Consistent with these results, we observed an increase in MT JNK phosphorylation (pJNK) in both conditions at an early time point (5 days of tumor induction) and at a late time point (8 days of tumor induction) (Fig. 4GH). Notably, the increase of JNK signaling is specific in the main segment region of the MT, since elevated levels of pJNK was only detected in the SC and PC region and not in the renal stem cell zone (SCZ) (Fig. 4I). Conversely, phosphorylation of ERK was not changed at late time points in Yki flies nor following PC-specific activation of PDGF/VEGF signaling (Fig. S4AB). Finally, knockdown of *Pvr* in PCs rescued elevated levels of pJNK (Fig. 4J). Altogether, these results indicate that the PDGF/VEGF signaling activates downstream JNK pathway in the MTs of Yki flies.

**Figure 4.**
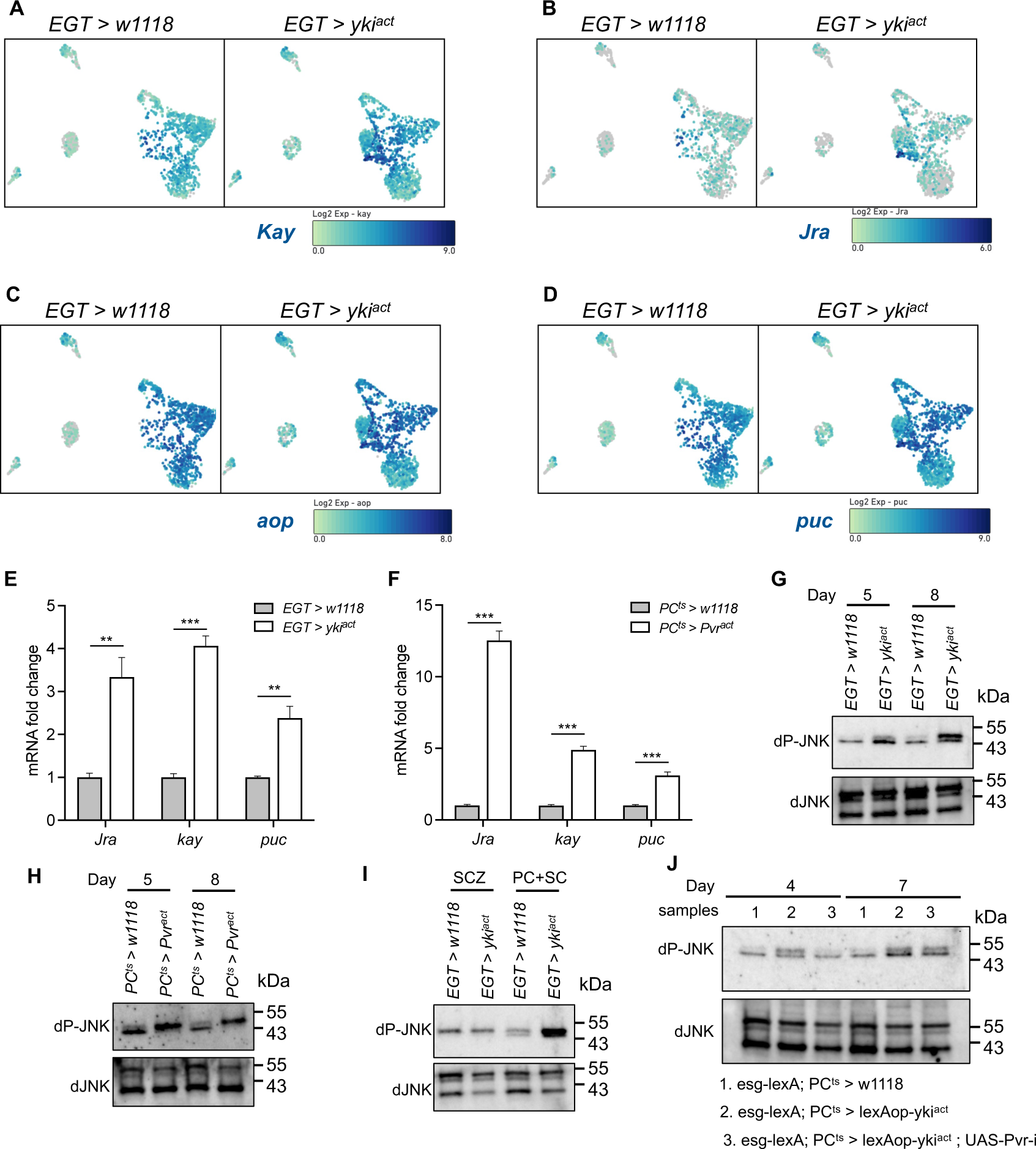
Increase in JNK/Jra signaling in principal cells in *EGT > yki^act^* and *PC^ts^ > Pvr^act^*flies. (A-D) UMAP showing the gene expression changes of JNK/Jra pathway genes in MT of *EGT > w1118* and *EGT > yki^act^* flies. (E and F) qPCR of *Jra*, *kay* and *puc* genes in MT of *EGT > yki^act^* and *PC^ts^ > Pvr^act^* flies. Transgenes were induced for 8 days. Data are presented as means ± SEM. *p < 0.05, **p < 0.01, ***p < 0.001 with student t-test. (G-J) Western blots showing the protein levels of JNK and phosphorylated JNK in MT of *EGT > w1118* and *EGT > yki^act^* (G) and *PC^ts^ > w1118* and *PC^ts^ > Pvr^act^* (I) flies. H shows the signal in the stem cell zone (SCZ) and principal cells plus stellate cells (PC+SC). J shows the rescue of p-JNK upon knockdown of *Pvr* in principal cells in yki^act^ flies. Transgenes were induced for 5 and 8 days in G and I; 8 days in H; 4 and 7 days in J.

### Function of principal cells depends on the transcription factor *Jra*

Next, we tested whether JNK signaling plays a role in tumor-induced renal dysfunction. While knockdown of *kay* in PCs did not rescue the MT defects in Yki flies, depletion of *Jra* was able to reverse bloating, kidney stone formation and uric acid elevation, without affecting the gut tumor (Fig. 5A-C). Importantly, inhibition of JNK signaling (by blocking *Bsk* via overexpression of *Bsk^DN^*) in *Pvr^act^* flies rescued bloating, lifespan, kidney stone formation, uric acid levels, and expression levels of genes involved in kidney function (Fig. 5D-I, Fig. S5). Previously, we reported that *Uro* is a target of *Jra*^26^, leading us to analyze the expression of *Uro* in PCs. *Uro* mRNA levels were up-regulated in MTs upon knockdown of *Jra* in PCs (Fig. 5J), suggesting that *Jra* inhibits *Uro* expression. Lower levels of *Uro* leads to insufficient oxidation of uric acid to 5-hydroxyisourate, which may contribute to the uric acid accumulation observed in Yki flies and in PCs with activated PDGF/VEGF signaling. To confirm this, we overexpressed *Uro* in PCs expressing *Pvr^act^*, which rescued whole-body levels of uric acid in these flies (Fig. 5K). Collectively, these observations indicate that PDGF/VEGF signaling regulates PCs function through the TF *Jra*.

**Figure 5.**
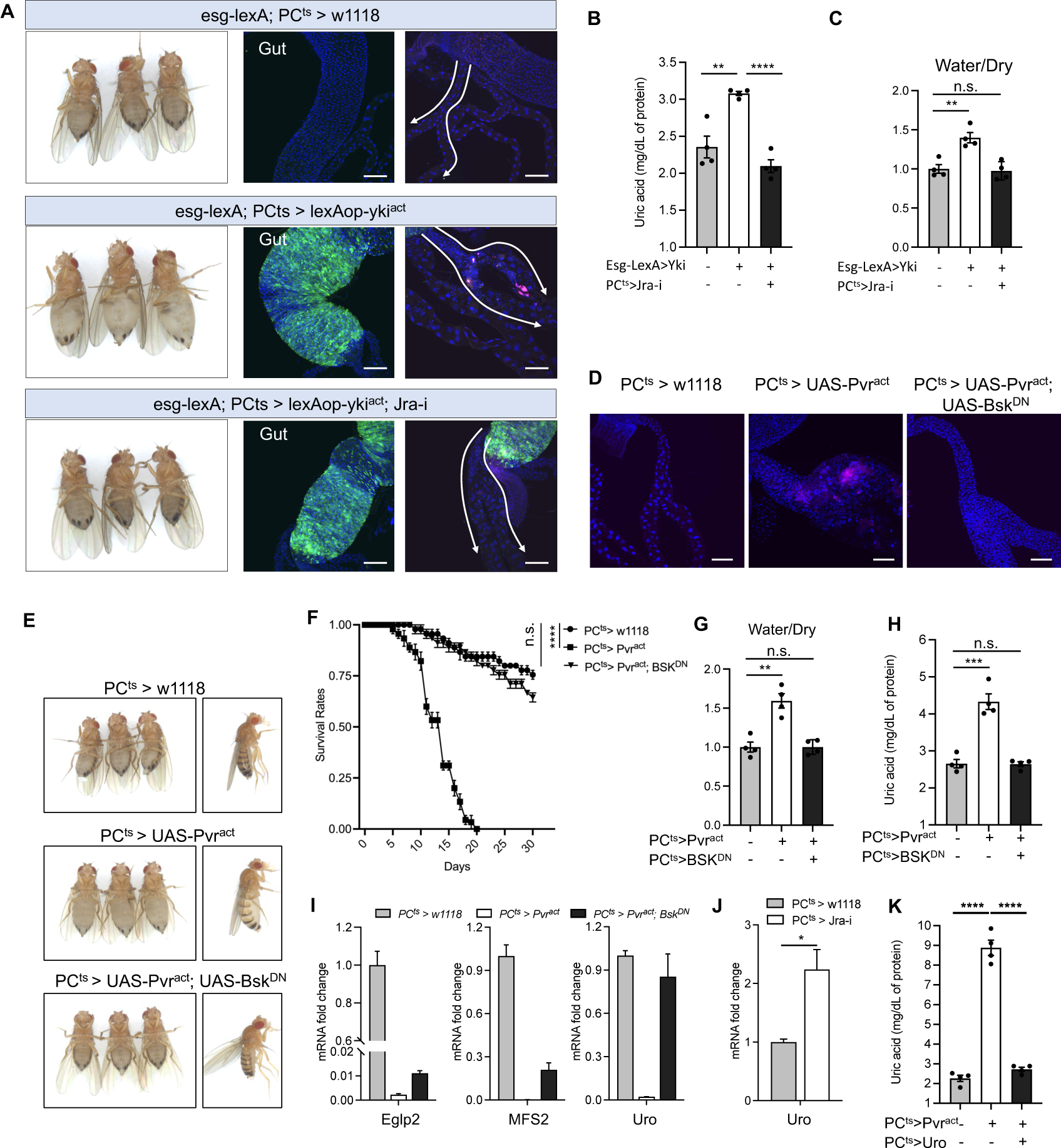
*Jra* targets *Uro* to control uric acid metabolism in MT. (A-C) Rescue of the bloating, kidney stone and uric acid phenotypes upon knocking down *Jra* in principal cells in yki^act^ flies. Transgenes were induced for 6 days. Rescue of (D) kidney stones, (E) bloating, (F) survival, (G) water/dry mass, (H) UA levels, and (I) mis-regulated gene expression (*Eglp2*, *MFS2*, and *Uro*) when *Pvr^act^* is co-expressed with *Bsk^DN^* in principal cells. (J) qPCR of *Uro* in the MT of *PC^ts^ > w1118* and *PC^ts^ > Jra-i* flies. Transgenes were induced for 8 days. (K) Rescue of uric acid level when *Pvr^act^* is co-expressed with *Uro* in principal cells. Data are presented as means ± SEM. *p < 0.05, **p < 0.01, ****p < 0.0001 with student t-test.

### Pvf1 from Yki tumors activates PVR signaling in renal PCs

Yki tumors secrete the Pvf1 ligand, activating PVR signaling pathway in peripheral tissues^15^. To determine whether tumor-derived Pvf1 activates PVR signaling pathway in MTs, we decreased *Pvf1* expression in the gut stem cells of Yki flies (*esg> yki^act^ + Pvf1-i*). As reported previously, inhibition of *Pvf1* in gut tumors rescues both bloating and the water/dry mass phenotype in Yki flies without affecting the tumor (Fig. 6AB)^15^. Strikingly, *Uro* expression and uric acid levels were decreased, and no kidney stones were observed in MTs (Fig. 6C-E). Importantly, inhibition of *Pvf1* in gut tumors rescued the expression of mis-regulated genes in the MTs of Yki flies (Fig. 6E, Fig.S6). Collectively, our data suggest that Pvf1 from gut tumor cells remotely activates PvR signaling in MTs, leading to renal dysregulation (Fig. 6F).

**Figure 6.**
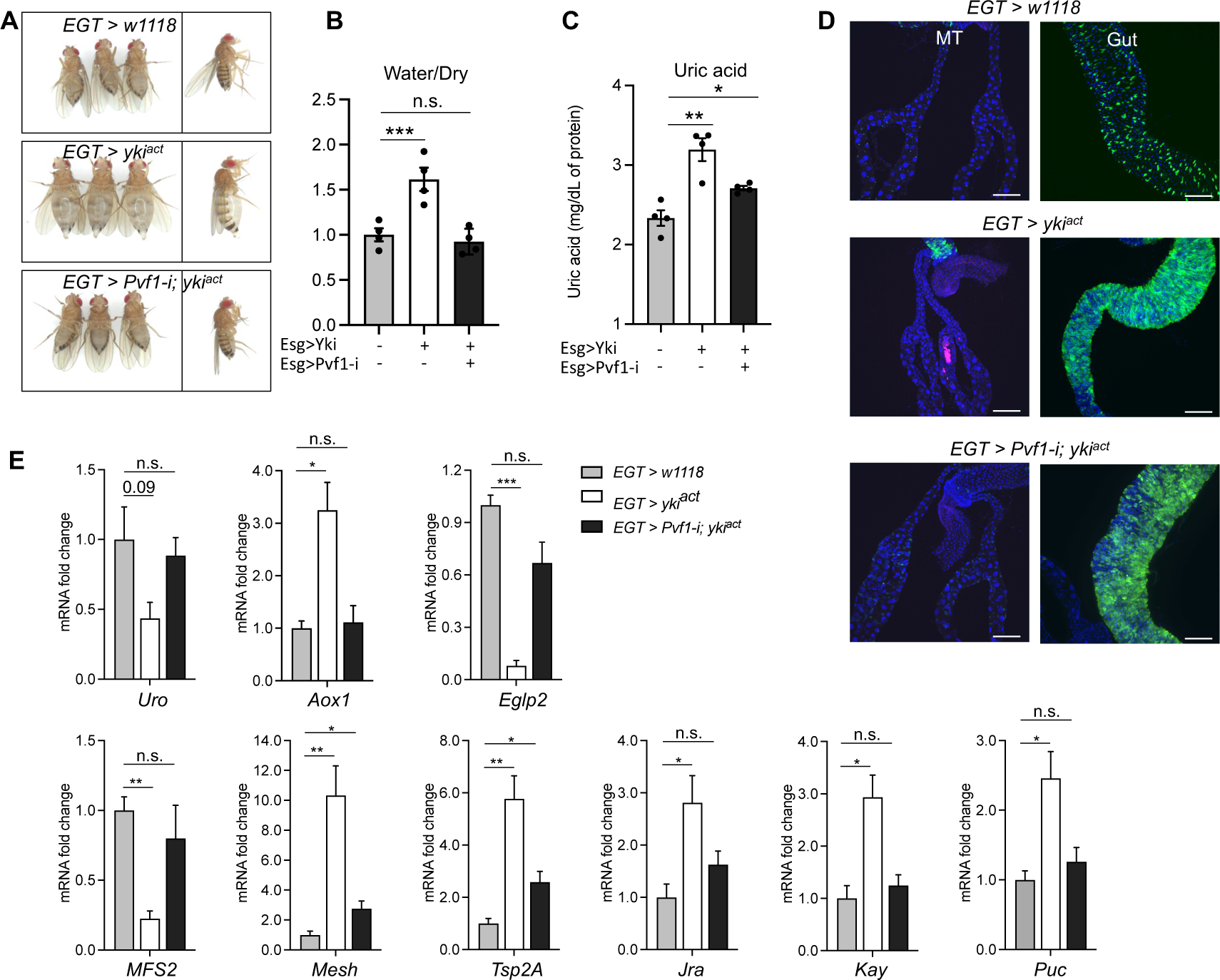
Knockdown of *Pvf1* in *EGT > yki^act^* flies repairs MT function. (A-D) Rescue of the bloating, uric acid, and kidney stone phenotypes upon knocking down *Pvf1* in gut stem cells in yki^act^ flies. Transgenes were induced for 8 days. (E) qPCR of kidney function and JNK pathway genes in the MT of *EGT > w1118*, *EGT > yki^act^* and *EGT > Pvf1-i; yki^act^* flies. Data are presented as means ± SEM. *p < 0.05, **p < 0.01, ***p < 0.001 with student t-test.

### Yki tumors hijack paracrine renal PDGF/VEGF signaling

In wildtype MTs, *Pvf1* is expressed in stellate cells (SCs), a small group of cells involved in fluid secretion, and *Pvr* is widely expressed in all PCs^17, 26^, suggesting that Pvf1 acts as a paracrine signal from SCs to PCs. The finding that Pvf1 derived from gut tumors acts as an endocrine hormone that activates PDGF/VEGF signaling in PCs suggests that tumor secreted Pvf1 may interfere with Pvf1 paracrine signaling in MTs. To characterize and compare the physiological roles of PDGF/VEGF signaling in wildtype and Yki flies, we performed snRNA-seq analysis of MTs of control flies (*PC^ts^ > w1118*) and flies with PDGF/VEGF signaling inhibition or activation in PCs (*PC^ts^ > Pvr^RN^* and *PC^ts^ > Pvr^act^*, respectively) (Fig. 7A). Inhibition of PDGF/VEGF signaling in PCs did not change the cell cluster composition of MTs (Fig. 7A). However, reduction of *Pvr* activity in PCs was associated with a wide range of elevated expression of ribosomal protein genes (Table S1), suggesting impaired ribosome biogenesis^35^. In addition, expression of many transporters including *CG7720*, *salt*, *Irk1*, *Zip48C*, and *Vha14-1* (Fig. 7B-F) was up-regulated (Table S2), suggesting that PDGF/VEGF signaling inhibits their expression in wildtype MTs. Despite these changes in gene expression, flies with *Pvr* depletion in PCs did not show any obvious phenotypes.

**Figure 7.**
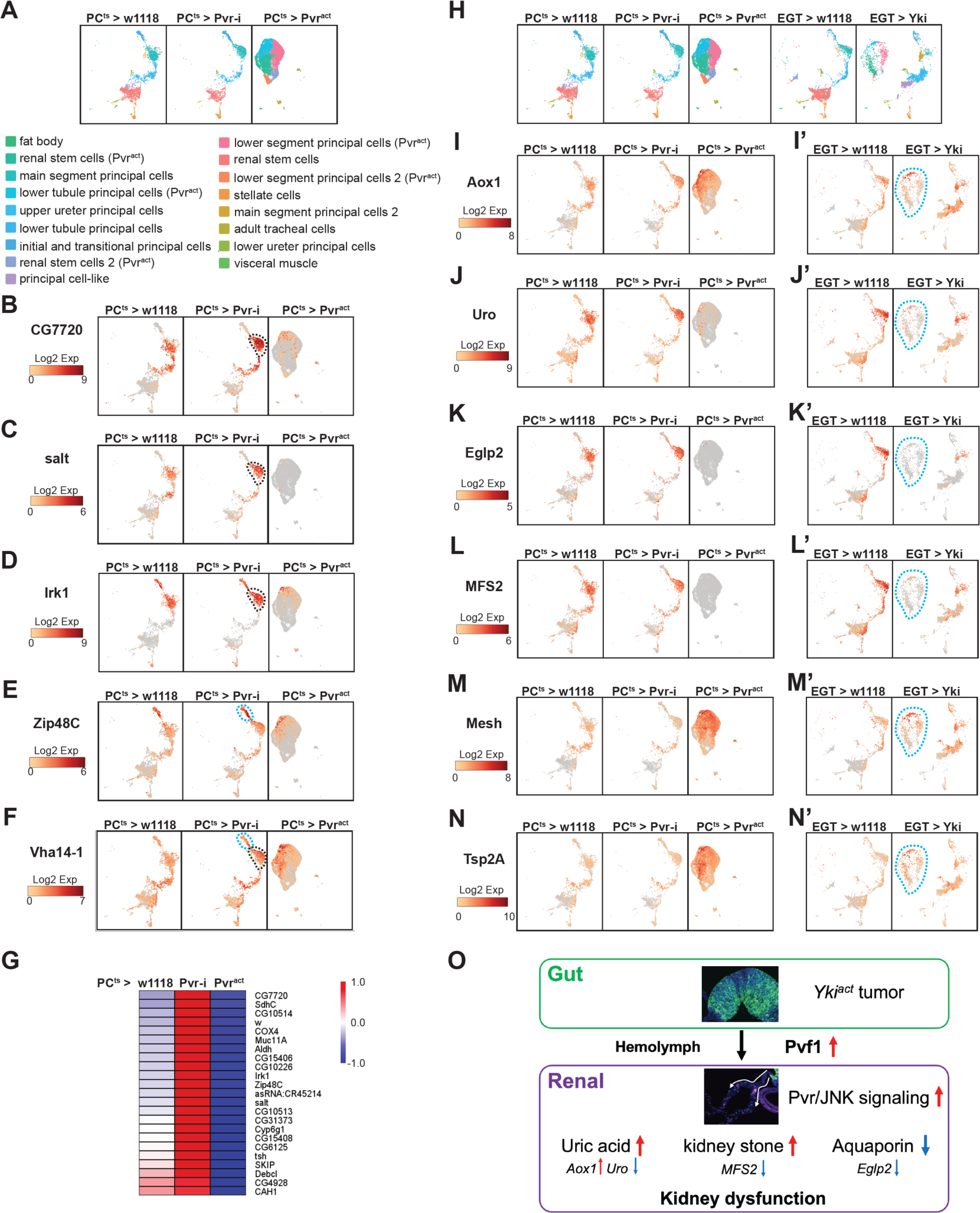
Gene expression in the MTs of *yki^act^* flies and flies with PDGF/VEGF signaling activation or inhibition. (A) UMAP indicating the MT cell types of flies with PDGF/VEGF signaling activation or inhibition. UMAP indicating (B) *CG7720*, (C) *salt*, (D) *Irk1*, (E) *Zip48C* and (F) *Vha14*-1 expression levels in MTs of control and flies with PDGF/VEGF signaling activation or inhibition. Dashed black circles indicate main segment principal cells and dashed cyan circles indicate initial and transitional principal cells. (G) Heatmaps showing expression changes of Pvr downstream target genes. (H) UMAP indicating the cell clustering of MT cells of control, Yki flies, and flies with PDGF/VEGF signaling activation or inhibition. UMAP indicating (I&I’) *Aox1*, (J&J’) *Uro*, (K&K’) *Eglp2*, (L&L’) *MFS2*, (M&M’) *Mesh*, and (N&N’) *Tsp2A* expression levels in MTs of control, Yki flies, and flies with PDGF/VEGF signaling activation or inhibition. Dashed cyan circles indicate *Pvr^act^* clusters in Yki fly MTs. (O) Model of *yki^act^* gut tumor induced renal dysfunction.

Unlike reduction of PDGF/VEGF signaling, activation of this pathway in PCs led to distinct MT cell clusters (referred to hereafter as *Pvr^act^* clusters) (Fig. 7A). In total, five *Pvr^act^* clusters were identified, including two renal stem cell-like clusters, two lower segment PC-like clusters, and a cluster related to lower tubule PCs (Fig. 7A). Presumably due to the high recovery rate of cells in *Pvr^act^* clusters, other MT cell clusters, such as SCs, were not detected (Fig. 7A). Considering that SCs represent a small cell population in WT, we decided to compare gene expression changes at the whole MT level among all three samples (*PC^ts^ > w1118, PC^ts^ > Pvr^RNAi^, and PC^ts^ > Pvr^act^*) (Table S2). Interestingly, 23 genes down-regulated in *Pvr^act^* MTs (Fig. 7G) were among the genes significantly up-regulated in *Pvr^RNAi^* MTs (Table S2), suggesting that these genes are downstream targets of the PDGF/VEGF pathway. Notably, among them are 9 transporter genes: *CG7720*, *CG15406*, *CG10226*, *Irk1*, *Zip48C*, *salt*, *CG15408*, *CG6125*, and *CG4928* (Fig. 7G, Table S2). In addition, 19 genes significantly down-regulated in *Pvr^RNAi^* MTs are up-regulated in *Pvr^act^* MTs, although no enrichment of gene groups were observed (Table S2). These data suggest that in wildtype MTs, SCs secrete Pvf1 to down-regulate the expression of transporter genes in PCs, possibly to regulate fluid secretion and ionic balance^17^.

Next, we compared the expression of the mis-regulated genes identified in the MTs of Yki flies with those of *PC^ts^ > w1118, PC^ts^ > Pvr^RNAi^,* and *PC^ts^ > Pvr^act^* MTs. Interestingly, *Aox1*, *Mesh*, and *Tsp2A* were up-regulated and *Uro*, *Eglp2*, *MFS2* were down-regulated, in both *Pvr^act^* MTs and the MTs of Yki flies (Fig. 7I-N, Fig. 2D, Table S2), indicating that activation of the PDGF/VEGF pathway in PCs in wildtype flies phenocopies the activation of this pathway in Yki flies. Next, we compared the single-cell transcriptome of MT cells in five conditions (*PC^ts^ > w1118*, *PC^ts^ > Pvr^RNAi^*, *PC^ts^ > Pvr^act^*, *esg> w1118*, and *esg> yki^act^*) (Fig. 7H). Interestingly, genes mis-regulated in the MTs of Yki flies were less changed than in MTs expressing *Pvr^act^* in PCs of wildtype flies (Fig. 7I-N, Fig. 7I’-N’), most likely reflecting that the Pvr pathway in Yki flies is not as active as in *PC^ts^ > Pvr^act^*. Consistent with this, MT cells of Yki flies appear to be a mixture of wildtype cells and *Pvr^act^*cells (Fig. 7H), and the mis-regulated genes identified in Yki flies displayed higher changes in *Pvr^act^* MTs (Fig.S7B, Table S2). Altogether, our results suggest that elevated Pvf1 derived from Yki gut tumors hijacks paracrine renal PDGF/VEGF signaling, leading to paraneoplastic defects in the renal system (Fig. 7O).

## Discussion

Paraneoplastic syndromes in cancer patients originate from the dysfunction of organs at a distance from tumors or their metastases. Despite the clinical relevance of paraneoplastic syndromes, relatively little is known about the mechanisms underlying their pathogenesis. In this study, we demonstrate that tumor-secreted Pvf1 activates the PDGF/VEGF/JNK pathway in the PC cells of MTs, leading to mis-expression of genes involved in aquaporin expression, kidney stone formation, cation transport, diuretic control, and uric acid metabolism. This pathological mis-regulation in gene expression leads to renal dysfunction and reduced viability, demonstrating a molecular mechanism of tumor-induced renal dysfunction.

Tumors hijack host signaling pathways to induce paraneoplastic syndromes. In the case of Yki flies, gut tumor-secreted Pvf1 targets muscle and adipose tissue, and activates the MEK/MAPK pathway, which induces muscle wasting and lipid loss that contribute to body wasting^15^. Here, we demonstrate that, in addition, tumor secreted Pvf1 activates the JNK pathway in MTs, leading to renal dysfunction. Abnormal activation of PDGF/VEGF/JNK signaling in PC cells of MTs leads to up-regulation of two key renal genes, *Aox1* and *Uro*, impairing uric acid excretion^21, 26, 27^ and resulting in an increase in uric acid levels. Further, high levels of uric acid and downregulation of *MFS2* induce formation of kidney stones^23, 28^. In addition, down-regulation of aquaporins such as *Eglp2* influence water transport^36^, which likely contributes to the excessive accumulation of body fluids observed in Yki flies. Interestingly, we observe a similar transcriptional up-regulation of *Aox1* and down-regulation of *Uro, MFS2* and *Eglp2*, as well as similar phenotypes, i.e., elevated levels of uric acid, kidney stone formation and bloating, in both Yki flies and flies with activated PDGF/VEGF signaling in PCs. Altogether, this suggests that the renal phenotypes observed in Yki flies are the result of activation of the PDGF/VEGF pathway in PCs. Notably, the paraneoplastic phenotypes in muscle and fat, and MTs, appear to result from activation of different signal transduction cascades, ERK in the case of muscle and fat, and JNK in MTs, reflecting tissue-specific differences in response to Pvf1.

In addition to identifying the role of Pvf1 as a paraneoplastic factor, our study implicates Pvf1 as a paracrine signal in WT MTs. Our recent study of cell composition of wildtype MTs indicated that *Pvf1* is expressed in SCs and that *Pvr* is enriched in PCs^26^. Strikingly, inhibition of *Pvr* in MT PCs led to an increase in expression of several transporter genes, suggesting that Pvf1 from SCs regulates fluid secretion and ionic balance in PCs^17^. Previously, we showed that Pvf1 from the muscles of adult flies regulates lipid synthesis in oenocytes^37^. It will be of interest to examine whether muscle derived Pvf1 can also regulate MT physiology.

Interestingly, loss of autocrine PDGF/VEGF signaling in wildtype MTs has no obvious phenotypes, although the expression of a number of transporter genes suggests a role of PVR signaling in regulating fluid secretion and ionic balance^17^. By contrast, elevated Pvf1 levels in Yki flies generate severe MT dysfunction and affect fly viability. These observations suggest that the physiological/pathological role of PDGF/VEGF depends on its activation levels. In mammals, VEGF signaling is important in formation and growth of blood vessels, and PDGF functions as a growth factor, which promotes proliferation and motility of mesenchymal and other cell types ^38, 39^. Both PDGF and VEGF signaling are important for normal development of the kidney, as evidenced from the kidney failure and death before or at birth of PDGF beta-receptor mutant mice^40^. In addition, abnormal glomeruli lacking capillary tufts and systemic edema phenotypes are observed in newborn mice with inhibition of VEGF signaling^41^. Interestingly, in mice and humans, PDGF and VEGF signaling mediate paracrine interactions among different types of kidney cells^42, 43^. In addition, increased activity of PDGF and VEGF signaling leads to kidney diseases such as renal fibrosis^44, 45^. These observations are reminiscent to the aberrant activation of paracrine PDGF/VEGF signaling leading to renal dysfunction observed in the MT of Yki flies. Further, both PDGF and VEGF signaling have reported roles in cancer progression^46, 47^; however, a role for PDGF/VEGF/JNK signaling in kidney physiology and paraneoplastic renal syndrome remains to be explored. Finally, defective kidney function has been reported in patients with cancer. While the cause of renal dysfunctions has been mainly attributed to cancer treatment, our study raises the possibility that tumor-secreted factors such as hormones and cytokines may contribute to the syndrome. Thus, it may be worth exploring whether kidney dysfunction in cancer patient involves a paraneoplastic role of PDGF/VEGF signaling.

## Materials and Methods

### Drosophila stocks

Fly husbandry and crosses were performed under a 12:12 hour light:dark photoperiod at 18°C, 25°C or 29°C, as indicated in each experiment. EGT (*esg-GAL4, UAS-GFP, tub-GAL80^TS^*), *esg-LexA* and *LexAop-yki^3SA^-GFP* are from the Perrimon lab stock collection.

The following strains were obtained from the Bloomington *Drosophila* Stock Center (BL), Vienna Drosophila Resource Center (V) and flyORF: *tsh-GAL4; tub-GAL80^TS^* (BL# 86330), *CG31272-GAL4* (BL# 76171), *UAS-yki^3SA^* (BL# 28817), *UAS-Pvr.lambda* (BL# 58428), *UAS-Pvr.lambda* (BL# 58496), *UAS-Pvr-RNAi* (BL# 37520), *UAS-upd3* is a kind gift from Dr. Frederic Geissmann, *UAS-InR.DN* (BL# 8253), *UAS-Jra-RNAi* (BL# 31595), *UAS-Pvf1-RNAi* (V#102699), and *UAS-Pvf1-3xHA* (#F002862). *UAS-Uro-127D01* fly containing the 127D01 nanotag (Xu et al., 2022) was generated in this study. *Uro* ORF was cloned from the pENTR-Uro (DGRC, #1656531) using the primers: ACTCTGAATAGGGAATTGGGAATTCCAAAATGTTTGCCACGCCCCTCAG & CAGATCAGAACTAGTTTGCTCTAGATTAATCCTCGCCTTTCCAGAAATCTTCAAAACTTCCTGAACCCAGGTGACTATTGATGTTCT, inserted into pWalium10 vector (DGRC, #1470). The vector was injected into attP line *y,w; P{nos-phiC31\int.NLS}X; P{CaryP}attP2* to generate the transgenic flies. Female flies are used in the analysis as they showed more significant and consistent bloating phenotype.

### *Drosophila* culture and drug treatment

Flies were raised on standard lab food (corn meal/agar medium). For conditional expression using tubGal80^ts^, flies were grown at 18°C until eclosion, maintained at 18°C for an additional 2d, and then shifted to 29°C to induce expression. The concentration of chemicals mixed in the fly food was based on previously published reports ^21, 23^. The Super CitriMax *Garcinia Cambogia* extract (Swanson) contains 60% hydroxycitric acid and was purchased from Amazon. Fly food was melted and mixed with a final concentration of 30 mg/ml *Garcinia Cambogia* extract ^23^. For NaOx feeding, a final concentration of 5 mg/ml sodium oxalate was mixed into melted fly food ^23^. The high purine food contains 20 mM adenine (Sigma-Aldrich, # A8626) and 20 mM guanine (Sigma-Aldrich, # G11950) ^21^.

### Lifespan analysis

For survival analysis, flies were collected within 24 hrs of eclosion (20 female flies and 5 males per vial) and cultured on normal lab fly food at 18°C and 60% humidity with 12 hrs on/off light cycle. After two days, matured and mated female flies were shifted to 29°C to induce gene expression. Flies were transferred to fresh vials every other day to keep the vials clean, and dead flies were counted every day.

### Uric acid, lipid and carbohydrate measurements

Uric acid was measured using the QuantiChrom^TM^ Uric Acid Assay Kit (Bioassay, DIUA-250). Whole fly carbohydrates and triglycerides were measured as described previously ^15, 19^. To prepare fly lysates for metabolic assays, we homogenized 4 flies from each group with 300 μl PBS supplemented with 0.2% Triton X-100 and heated at 70°C for 10 min. The supernatant was collected after centrifugation at 3000g for 1 min at 4°C. 10 μl of supernatant was used for protein quantification using Bradford Reagent (Sigma, B6916-500ML). Whole-body uric acid levels were measured from 5 μl of supernatant at 22°C for 30 min using uric acid assay kit following the manufacturer’s protocol. Whole-body trehalose levels were measured from 10 μl of supernatant treated with 0.2 μl trehalase (Megazyme; E-TREH) at 37°C for 30 min using glucose assay reagent (Megazyme; K-GLUC) following the manufacturer’s protocol. Whole-body glycogen levels were determined from 10 μl of supernatant preincubated with 1 μl amyloglucosidase (Sigma-Aldrich; A7420) at 37°C for 30 min using glucose assay reagent (Megazyme; K-GLUC). We subtracted the amount of free glucose from the measurement and then normalized the subtracted values to protein levels in the supernatant. To measure whole body triglycerides, we processed 10 μl of supernatant using a Serum Triglyceride Determination kit (Sigma, TR0100). We subtracted the amount of free glycerol from the measurement and then normalized the subtracted values to protein levels.

### Single nucleus isolation and sequencing

90 pairs of *Drosophila* MTs were dissociated for each sample and single nuclei were prepared as previously described ^26^ with a few modifications. MT were dissected under a microscope from *EGT>w1118* and *EGT > yki^act^* female adult flies after incubation at 29 °C for 8 days. Ten flies at a time were dissected and samples immediately transferred into 1.7 ml EP tube with Schneider’s medium on ice to avoid exposing the tissues to room temperature for a long period of time. Once 30 flies were dissected, EP tubes were sealed with parafilm and transferred into liquid nitrogen to quickly freeze the sample and storing samples in 100 ul Schneider’s medium at -80°C for long-term. After thawing, samples were spined down using a bench top spinner, medium was discarded, and 1000 ul homogenization butter was added. Nuclei were released by homogenizing a sample on ice with a 1 ml dounce homogenizer. The homogenized sample was then filtered through a cell strainer (35 um) and 40 um Flowmi. After centrifugation for 10 min at 1000 g at 4°C, nuclei were resuspended in PBS/0.5% BSA with RNase inhibitor. Before FACS sorting, samples were filtered again using 40 um Flowmi and nuclei were stained with Hoechst 33342. Ten thousand nuclei per sample were collected by FACS and loaded into a Chromium Controller (10X Genomics, PN-120223) on a Chromium Single Cell B Chip (10X Genomics, PN-120262), and processed to generate single cell gel beads in emulsion (GEM) according to the manufacturer’s protocol (10X Genomics, CG000183). The library was generated using the Chromium Single Cell 3′ Reagent Kits v3.1 (10X Genomics, PN-1000121) and Chromium i7 Multiplex Kit (10X Genomics, PN-120262) according to the manufacturer’s manual. Quality control for the constructed libraries was performed by Agilent Bioanalyzer High Sensitivity DNA kit (Agilent Technologies, 5067-4626). Quantification analysis was performed by Illumina Library Quantification Kit (KAPA Biosystems, KK4824). The library was sequenced on an Illumina NovaSeq system.

### snRNAseq dataset processing

The count matrices for each sample were generated using Cellranger count 7.0.0 under default setting and then imported into Seurat (4.3.0). All count data were normalized and scaled before batch corrected using Harmony (0.1.0) and subsequently reduced to 2-D Uniform Manifold Approximation and Projection (UMAP). With the corrected Harmony embeddings, each cell was connected with its closest 20 neighbors and assigned to clusters by Louvain algorithm under resolution=0.4. Differentially expressed genes for each cluster were computed with Wilcox Rank Sum test.

### Quantitative RT-PCR

Total RNA was extracted from 30 MT pairs per genotype from female flies per genotype using NucleoSpin RNA kit (Fisher Scientific, # NC9581114). cDNAs were synthesized using the iScript cDNA synthesis kit (Bio-Rad, #1708896) in a 20 μl reaction mixture containing 500 ng total RNA. Quantitative real-time RT-PCR (RT-qPCR) assays were performed using iQ SYBR Green Supermix (Bio-Rad, #1708880) on a CFX96 Real-Time PCR Detection System (Bio-Rad). RT-qPCR reactions were carried out with gene-specific primers (Table S3). A 5-fold serial dilution of pooled cDNA was used as the template for standard curves. Quantitative mRNA measurements were performed in three independent biological replicates and three mechanical replicates, and data were normalized to the amount of *Dmrp49* mRNA.

### Immunostaining and confocal microscopy

*Drosophila* MTs connected to guts were dissected from adult females and fixed in 4% paraformaldehyde in phosphate-buffered saline (PBS) at room temperature for 1 hour, incubated for 1 hour in Blocking Buffer (5% normal donkey serum, 0.3% Triton X-100, 0.1% bovine serum albumin (BSA) in PBS), and stained with primary antibodies overnight at 4°C in PBST (0.3% Triton X-100, 0.1% BSA in PBS). Primary antibodies were mouse anti-GFP (Invitrogen, A11120; 1:300) and mouse anti-discs-large (DSHB, 4F3,1:50). After primary antibody incubation, the tissues were washed 4 times with PBST, stained with 4′,6-diamidino-2-phenylindole (DAPI) (1:2000 dilution) and Alexa Fluor-conjugated donkey-anti-mouse (Molecular Probes, 1:1000), in PBST at room temperature for 2 hours, washed 4 times with PBST, and mounted in Vectashield medium.

All images presented in this study are confocal images captured with a Nikon Ti2 Spinning Disk confocal microscope. Z-stacks of 15-20 images covering one layer of the epithelium from the apical to the basal side were obtained, adjusted, and assembled using NIH Fiji (ImageJ), and shown as a maximum projection.

### Western blots

20 MTs pairs per genotype from female flies were dissected in PBS, placed in 30μl 2xSDS sample buffer (Thermo Scientific, #39001) containing 5% 2-Mercaptoethanol at 100°C for 10 minutes, ran 4ul on a 4%-20% polyacrylamide gel (Bio-Rad, #4561096), and transferred to an Immobilon-P polyvinylidene fluoride (PVDF) membrane (Millipore, IPVH00010). Membranes were blocked by 5% skim milk in 1x Tris-buffered saline (TBS) containing 0.1% Tween-20 (TBST) at room temperature for 30 minutes. The following primary antibodies were used: mouse anti-tubulin (Sigma, T5168, 1:10,000), rabbit anti-JNK Antibody (D-2) (Santa Cruz, sc-7345, 1:1000), rabbit phospho-JNK (Cell Signaling, 4668T, 1:1000), rabbit anti-ERK Antibody (Cell Signaling, 4695, 1:1000), rabbit phospho-ERK (Cell Signaling, 4370, 1:1000). After washing with TBST, signals were detected with enhanced chemiluminescence (ECL) reagents (Amersham, RPN2209; Pierce, #34095). Western blot images were acquired by Bio-Rad ChemiDoc MP.

**ACKNOWLEDGMENTS.** We thank the assistance provided by the Microscopy Resources on the North Quad (MicRoN) core and Biopolymers Facility at Harvard Medical School. We thank Dr. Frederic Geissmann for sharing fly stocks, Mikhail Kouzminov for help with snRNA-sequencing data analysis, and Stephanie Mohr for comments on the manuscript. J.X. is supported by the start-up funding from Shanghai Institute of Plant Physiology and Ecology/Center for Excellence in Molecular Plant Sciences, Chinese Academy of Sciences. Y.L is supported by Sigrid Jusélius Foundation and Finnish Cultural Foundation (Suomen Kulttuurirahasto). N.P. is an investigator of Howard Hughes Medical Institute.

This article is subject to HHMI’s Open Access to Publications policy. HHMI lab heads have previously granted a nonexclusive CC BY 4.0 license to the public and a sublicensable license to HHMI in their research articles. Pursuant to those licenses, the author-accepted manuscript of this article can be made freely available under a CC BY 4.0 license immediately upon publication.

**Author contributions:** N.P., Y.L. and J.X. conceptualized and designed the experiments. J.X. and Y. L. performed most of experiments. J.S.S.L. performed some fly work. Weihang Chen, Aram Comjean, Yanhui Hu performed the bioinformatics analyses. J.X., Y.L., and N.P. analyzed the data. J.X. wrote the first draft of the paper. Y. L., J.S.S.L. and N.P. edited the paper. All authors discussed the results and commented on the paper.

## Supplementary data

**Figure S1-related to Fig. 1.**
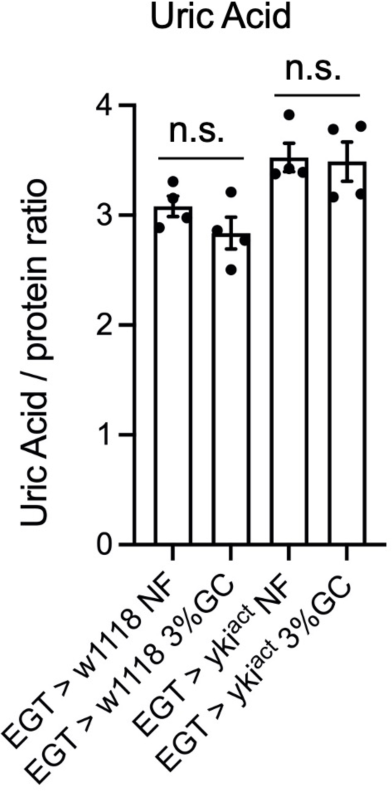
(A) Whole-body levels of uric acid in *EGT > w1118* and *EGT > yki^act^* flies at 8 days. Flies were fed either normal food or food with 3% *Garcinia cambogia* (GC). Data are presented as means ± SEM. n.s. means no significant with student t-test.

**Figure S2-related to Fig. 2.**
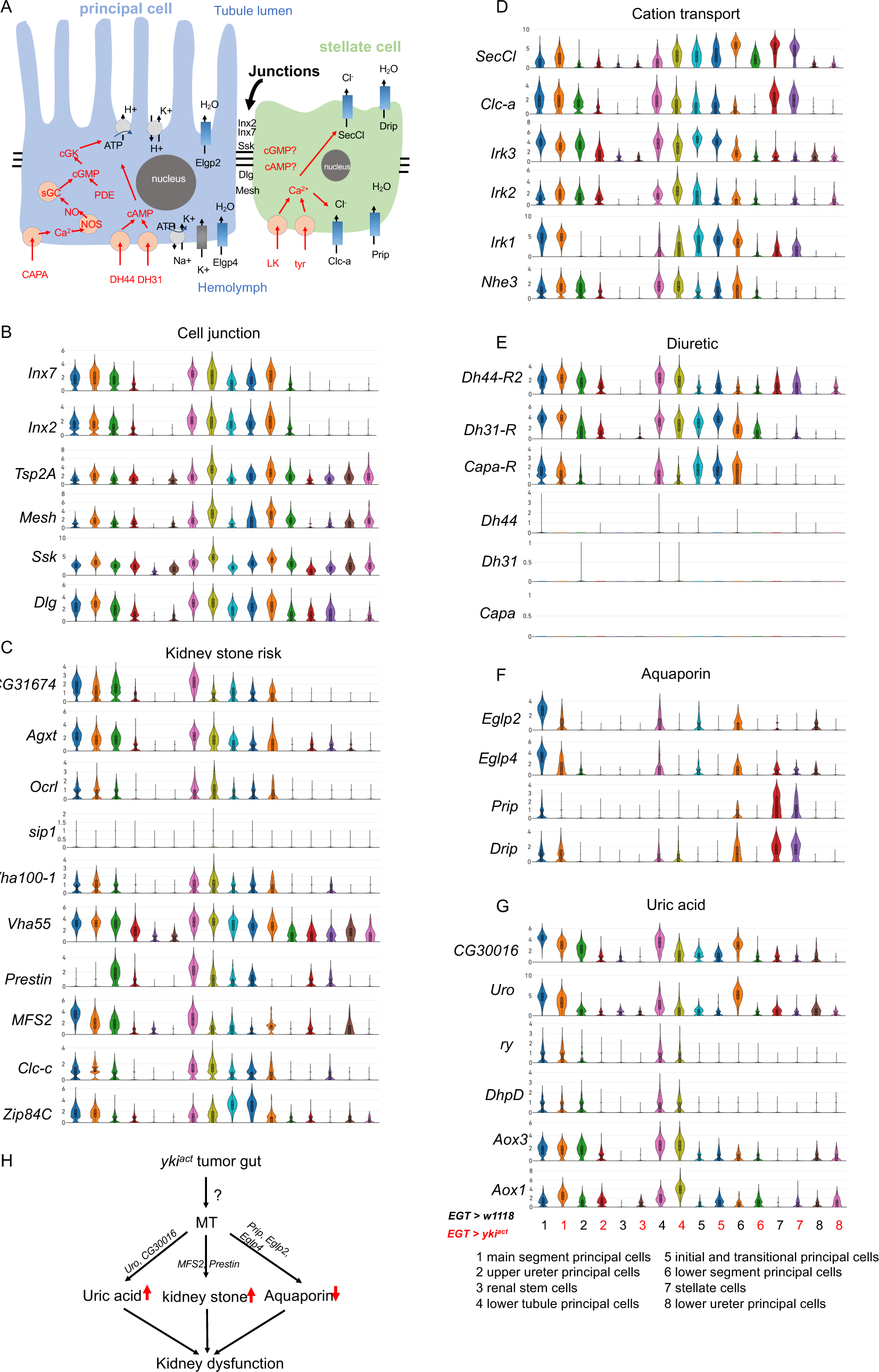
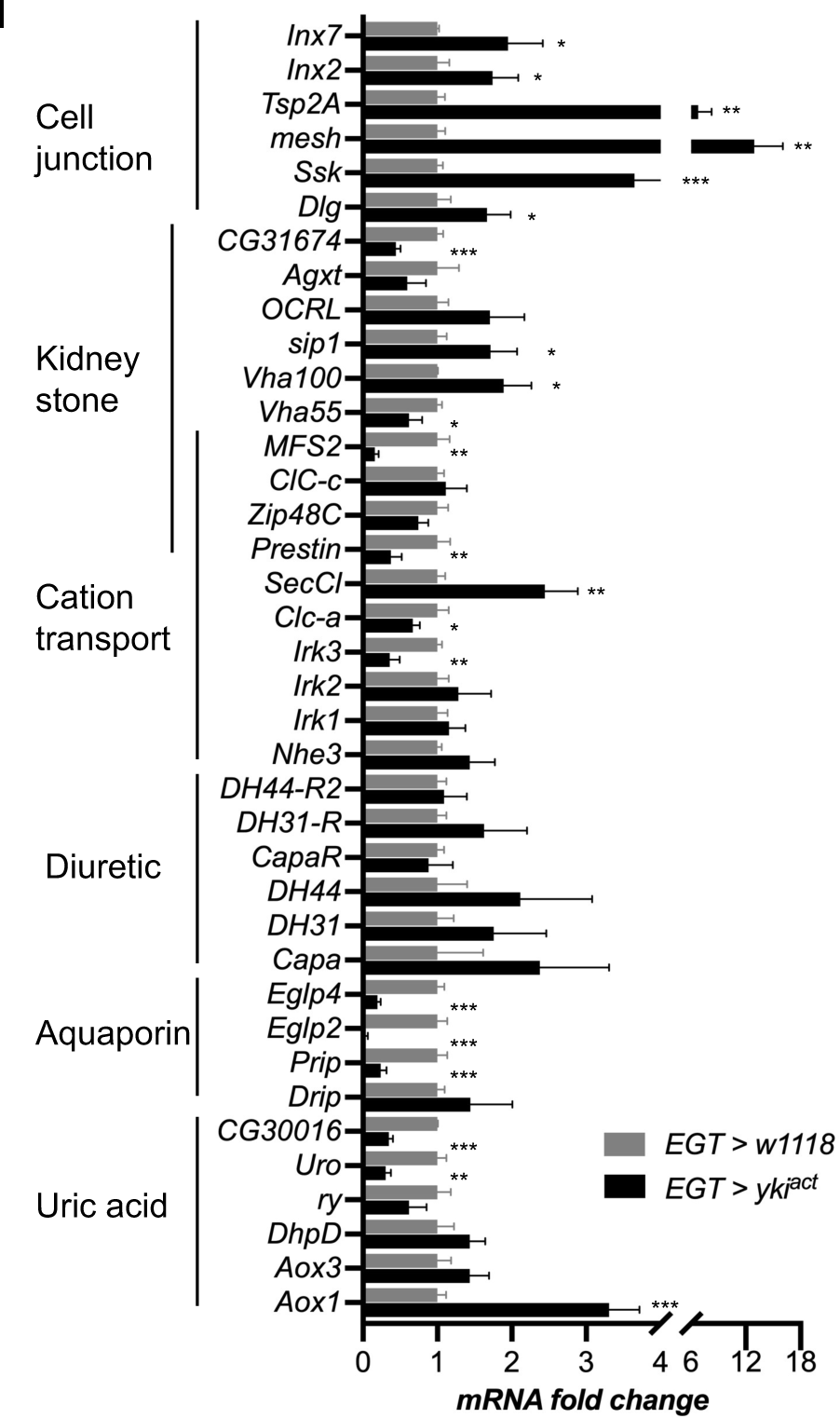
Changes in the expression of genes involved in MT function. (A) Function of principal and stellate cells in MT. Adapted from Xu et al., 2022. (B-G) Violin plots showing the expression of genes involved in kidney function (cell junction, kidney stone risk, cation transport, diuretic, aquaporin, uric acid) in the different cell clusters between *EGT > w1118* and *EGT > yki^act^*. (H) Model of renal dysfunction based on the expression of genes involved in uric acid synthesis, kidney stone formation and water transport in *yki^act^* flies. (I) qPCR analysis indicating the changes in gene expression levels in the MT of *EGT > w1118* and *EGT > yki^act^*. Transgene expression was induced for 8 days. Data are presented as means ± SEM. *p < 0.05, **p < 0.01, ***p < 0.001 with student t-test.

**Figure S3-related to Fig. 3.**
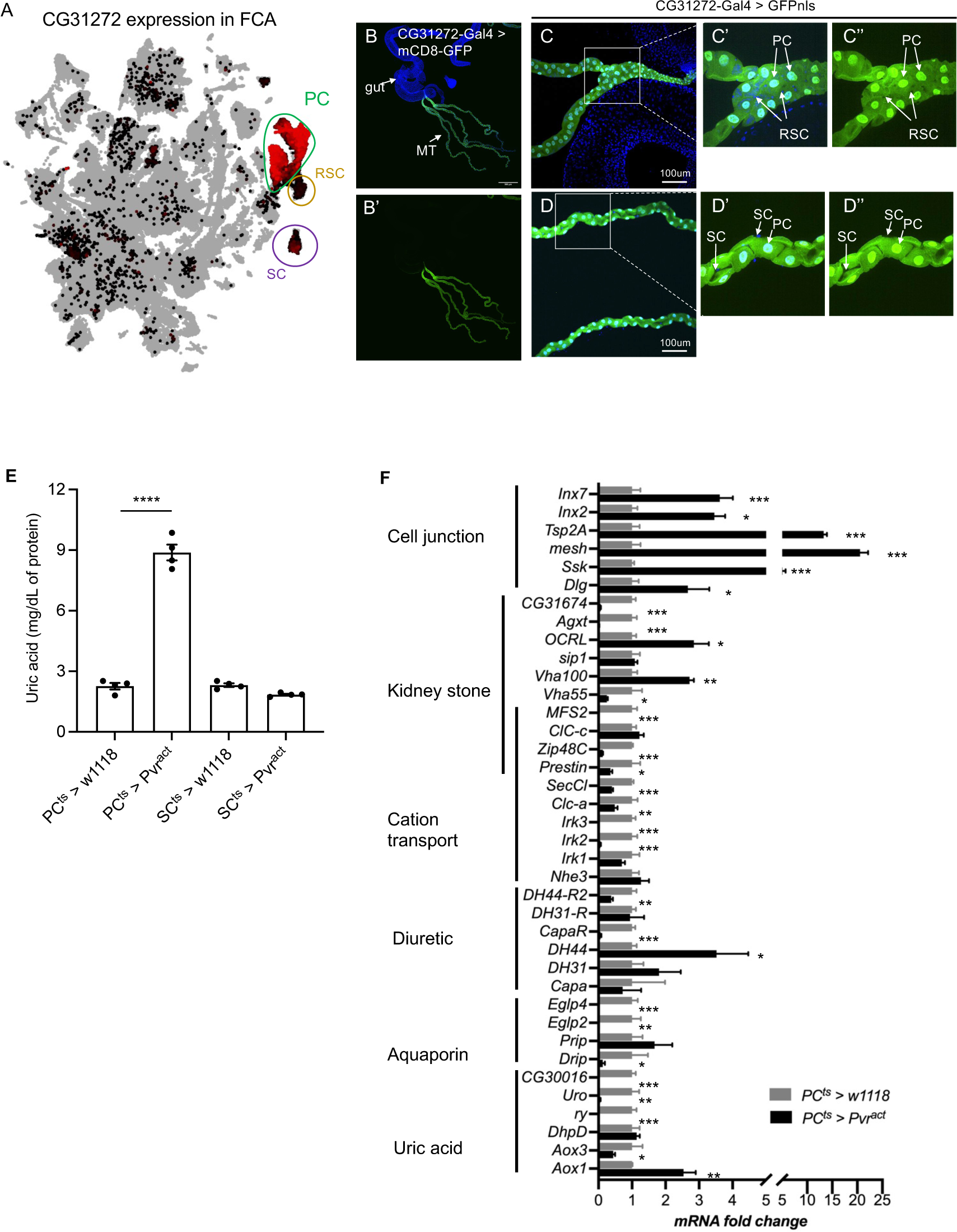
(A) FCA cell clusters with red indicates CG31272 expression. CG31272 is highly expressed in principal cells (PCs), and with a much lower expression level in renal stem cells (RSC) and stellate cells (SC) (data obtained from the FCA dataset; Li et al., 2022). (B-D) Characterization of a new Gal4 driver for MT principal cells (PCs). (E) Whole-body UA level in the *PC^ts^ > Pvr^act^, Pvf1, and SC^ts^ > Pvr^act^, Pvf1*, and control flies (n=4). (F) qPCR analysis indicating the changes in gene expression levels in the MT of *PC^ts^ > w1118* and *PC^ts^ > Pvr^act^*. Transgene expression was induced for 8 days. Data are presented as means ± SEM. *p < 0.05, **p < 0.01, ***p < 0.001 with student t-test.

**Figure S4-related to Fig. 4.**
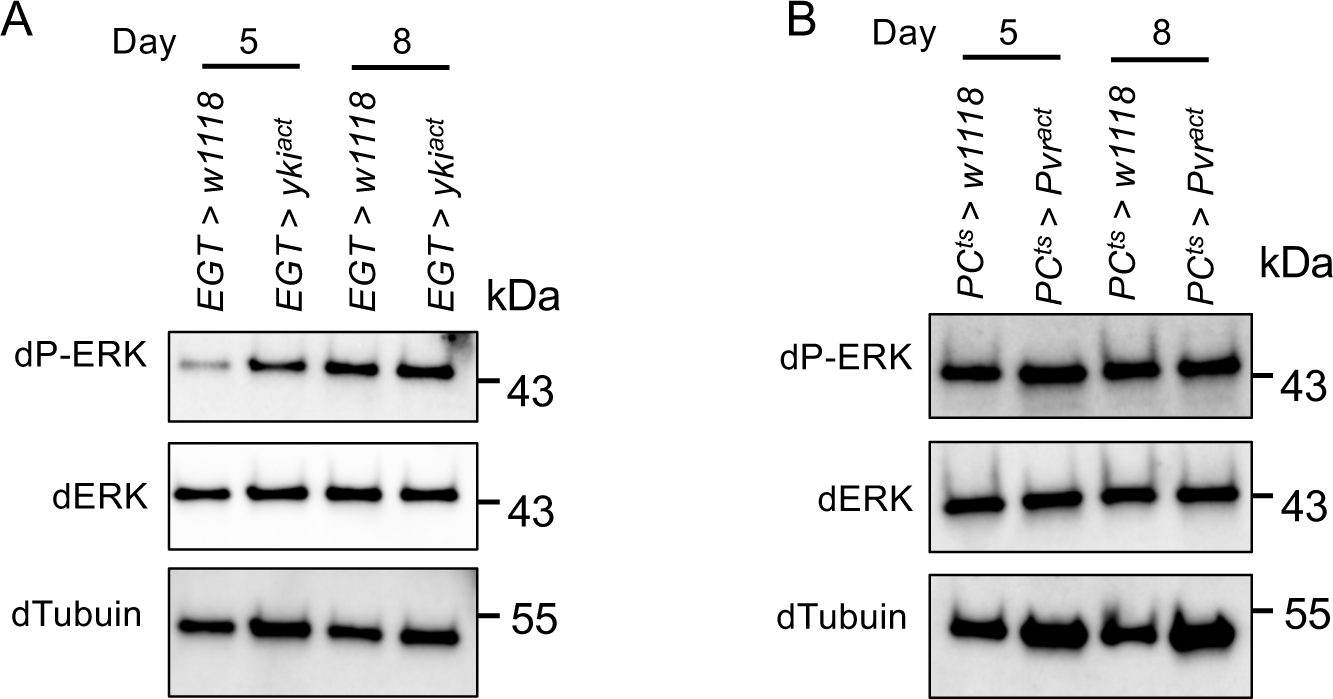
Western blots indicating the protein level of ERK and phosphorylated ERK in the MTs of **(A)** *EGT > w1118* and *EGT > yki^act^* and **(B)** *PC^ts^ > w1118* and *PC^ts^ > Pvr^act^*flies. Transgenes were induced for 8 days.

**Figure S5-related to Fig. 5.**
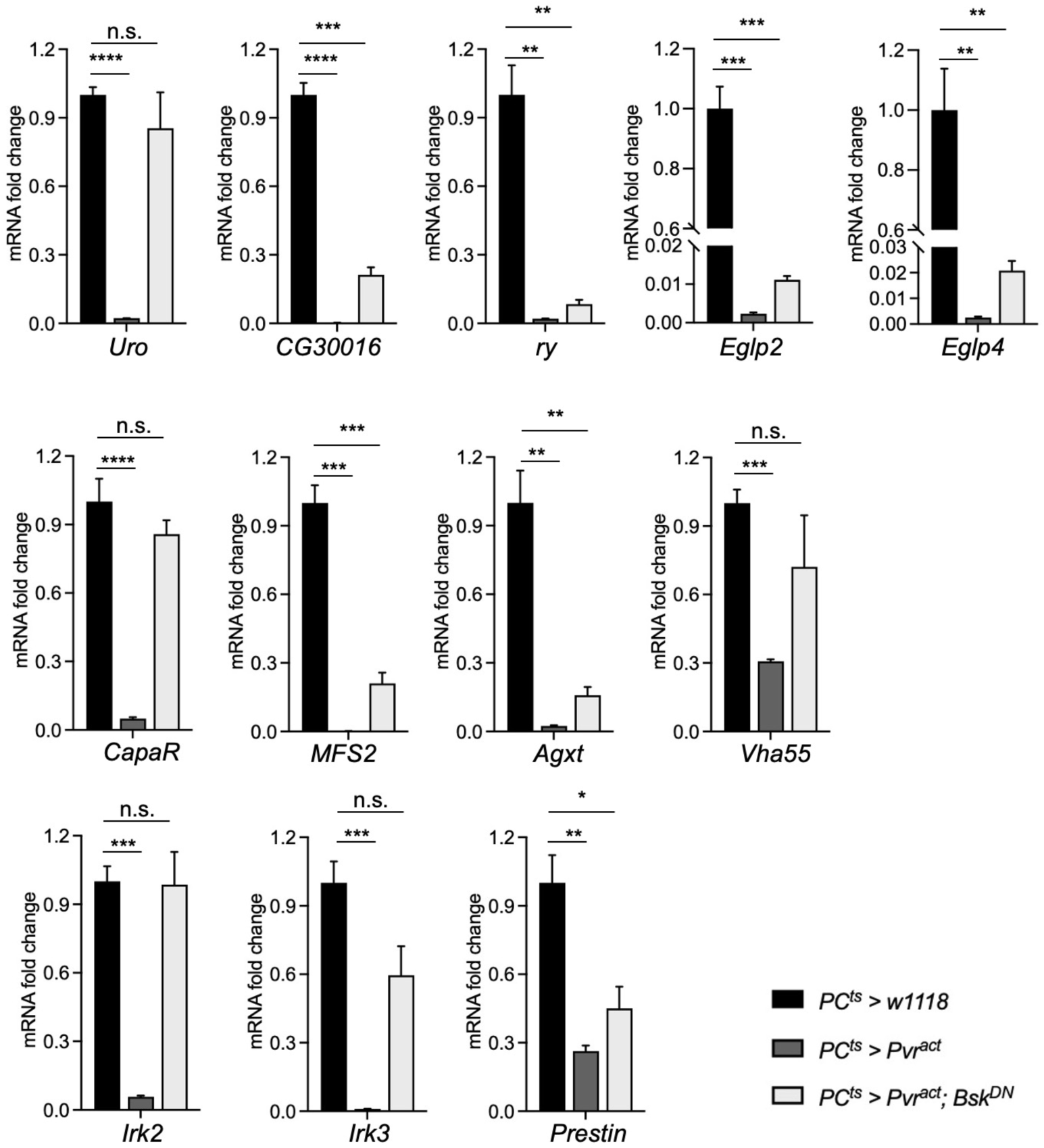
Expression of genes involved in kidney function. qPCR results showing changes of gene expression level in the MTs of *PC^ts^ > Pvr^act^*, *PC^ts^ > Pvr^act^; Bsk^DN^* and control flies.

**Figure S6-related to Fig. 6.**
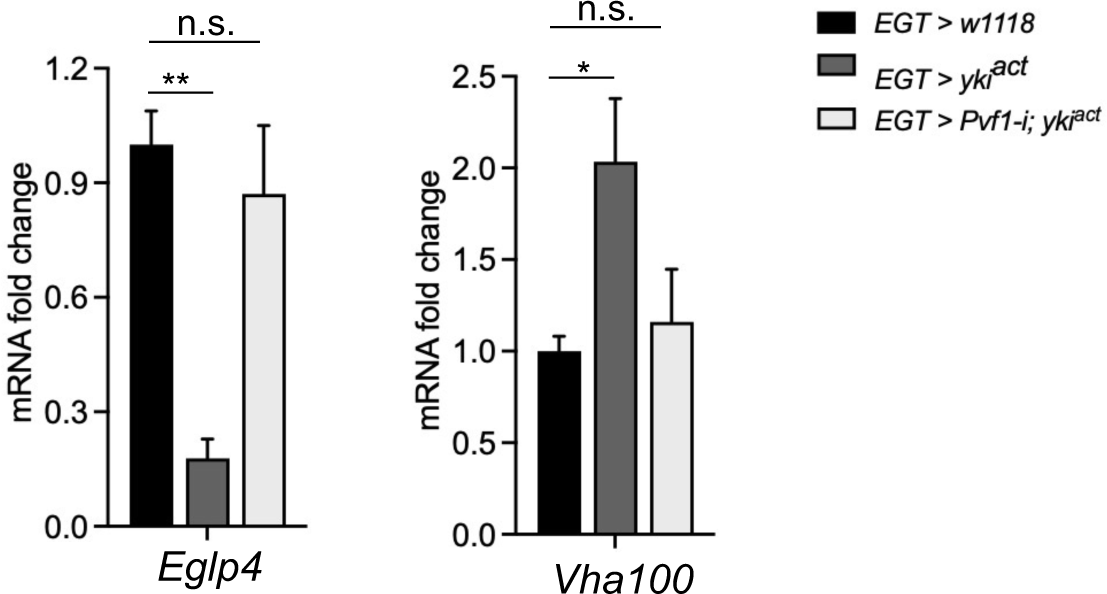
Expression of genes involved in kidney function. qPCR results showing changes in *Eglp4* and *Vha10*0 expression levels in the MTs of Yki flies with and without ISC depletion of Pvf1 and control flies.

**Figure S7-related to Fig. 7.**
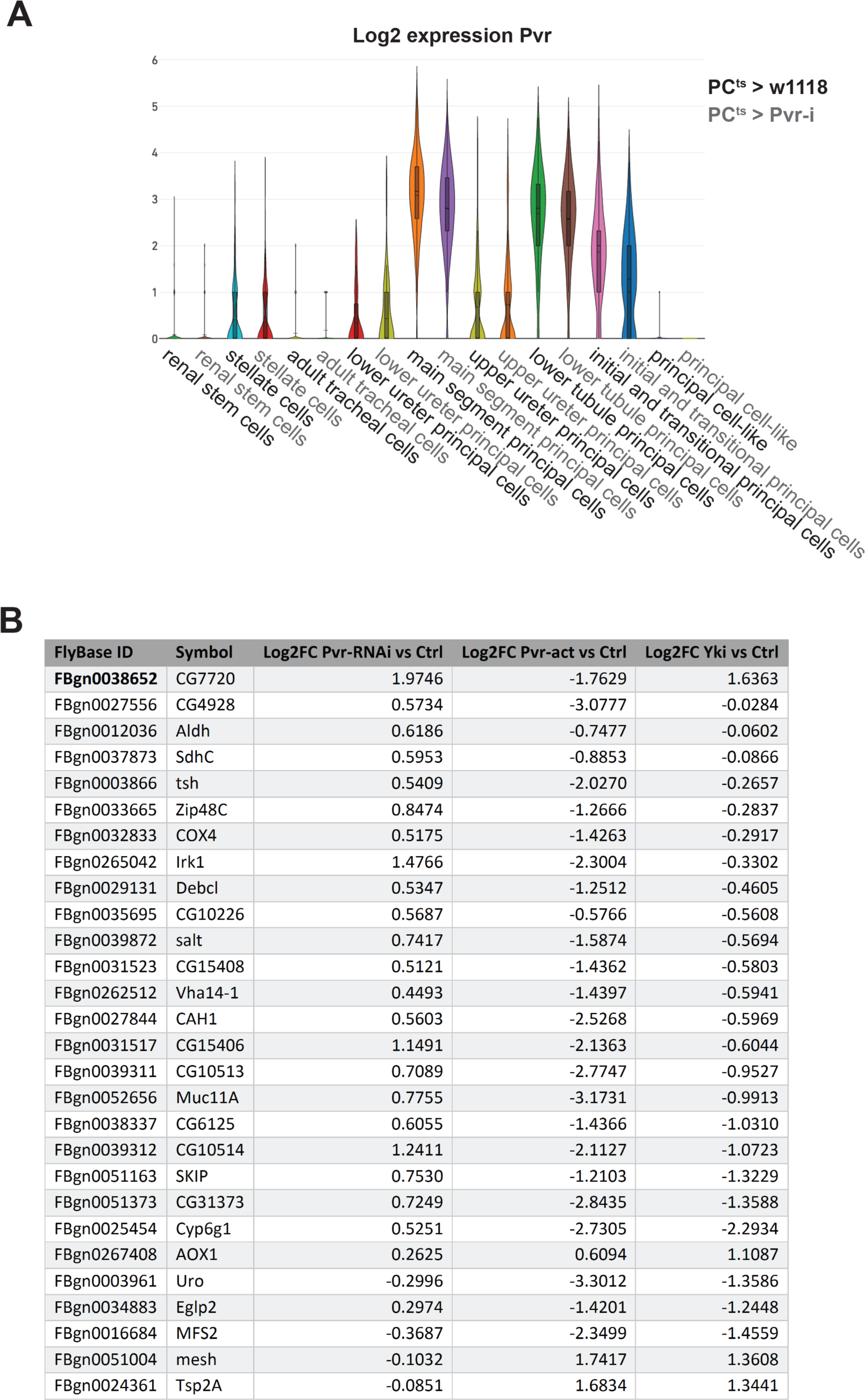
snRNAseq of flies with renal PDGF/VEGF signaling activation or inhibition. (A) Expression levels of *Pvr* in *PC^ts^ > w1118* and *PC^ts^ > Pvr^RNAi^* MTs visualized by violin plots. (B) Expression alternations of selected genes, full list sees Table S2.

